# Hoxb5+ fetal liver hematopoietic stem cells establish lifelong hematopoiesis and exhibit enhanced ITGA4-dependent engraftment

**DOI:** 10.64898/2026.06.30.731692

**Authors:** Allison Banuelos, Michelle Baez, Leyla Yılmaz, Elle Koren-Sedova, Allison Zhang, Monika Zukowska, Nicole Womack-Gambrel, Mary Moffitt, Andrew T. Burden, Victoria L. Mascetti, Ruby Honjol, Jinyi Xiang, Rahul Sinha, Irving L. Weissman

## Abstract

Adult long-term hematopoietic stem cells (LT-HSCs) are classically defined by self-renewal, multilineage regenerative capacity, and relative quiescence, but how and when lifelong LT-HSCs are established during development remains unclear. Here, we demonstrate that *Hoxb5*⁺ fetal liver HSCs exhibit bona fide LT-HSC activity, including long-term multilineage reconstitution and serial transplantation capacity, whereas *Hoxb5*⁻ fetal liver HSCs display limited regenerative potential. Embryonic lineage tracing further demonstrates that E14.5 *Hoxb5*-expressing hematopoietic cells contribute broadly to adult hematopoiesis, including the adult HSC compartment, and give rise to functional adult LT-HSCs. Across developmental stages, single-cell transcriptional profiling revealed that fetal *Hoxb5*⁺ HSCs remain highly proliferative while maintaining canonical LT-HSC transcriptional programs and superior repopulating activity relative to predominantly quiescent adult *Hoxb5*⁺ HSCs. Fetal *Hoxb5*⁺ HSCs also exhibited elevated ITGA4-mediated adhesion programs, and disruption of the ITGA4–VCAM1 axis impaired engraftment following transplantation. Together, these findings establish a developmental continuum linking fetal and adult LT-HSCs and identify enhanced ITGA4-mediated adhesion as a defining feature of fetal LT-HSCs.

## Introduction

Hematopoietic stem cells (HSCs) sustain lifelong blood and immune cell production through their unique capacity for self-renewal and multilineage differentiation^1–4^. During embryogenesis, HSCs arise through a series of developmental transitions before establishing the adult hematopoietic system^5–8^. Following their emergence, the fetal liver serves as the major site of HSC expansion during mid-gestation, providing a specialized niche that supports rapid proliferation, maturation, and subsequent colonization of the bone marrow^9,10^. This developmental window is critical for establishing the long-term HSC (LT-HSC) pool that maintains hematopoiesis throughout life^11,12^.

Although fetal liver HSCs undergo extensive expansion during development and are thought to contribute substantially to the adult hematopoietic system, the specific fetal HSC populations that persist into adulthood and sustain lifelong blood production remain incompletely defined^13–16^. The fetal liver contains phenotypically and functionally heterogeneous HSC populations, yet it remains unclear which fetal HSCs give rise to the adult LT-HSC compartment and whether developmental LT-HSC identity is established during fetal life or acquired later following bone marrow colonization^17–19^.

Adult LT-HSCs are classically characterized by long-term self-renewal, multilineage regenerative capacity, and relative quiescence^20–22^. In contrast, fetal liver HSCs exist within a highly proliferative developmental environment and undergo extensive expansion during mid-gestation^9,10,23^. This distinction raises fundamental questions regarding the developmental origins of lifelong hematopoietic stem cell function. Specifically, it remains unclear whether durable LT-HSC activity is acquired only following the establishment of adult quiescence or is already present within actively cycling fetal HSC populations. Although fetal HSCs are known to exhibit robust engraftment potential^9,20^, the relationship between proliferation, self-renewal, and the establishment of lifelong LT-HSC identity remains incompletely understood.

*Hoxb5* was previously identified as a highly specific marker of adult LT-HSCs^24^, raising the possibility that *Hoxb5* expression may define a developmentally conserved population of definitive LT-HSCs. Whether Hoxb5-expressing fetal liver HSCs represent functional LT-HSCs that contribute to the adult hematopoietic system, and how their properties relate to those of adult LT-HSCs, remains unknown.

Here, we identify Hoxb5⁺ fetal liver HSCs as a population with bona fide LT-HSC activity, including durable multilineage reconstitution and serial self-renewal. Using embryonic lineage tracing, we demonstrate that Hoxb5-expressing fetal hematopoietic cells contribute to the adult HSC compartment and give rise to functional adult LT-HSCs with long-term regenerative capacity. We further show that fetal Hoxb5⁺ HSCs retain robust stem cell function despite extensive proliferative activity and utilize ITGA4-dependent adhesive interactions that support efficient engraftment. Together, these findings establish a developmental continuum linking fetal and adult LT-HSCs and identify *Hoxb5* as a marker of definitive mouse LT-HSCs across ontogeny.

## Results

### *Hoxb5+* FL HSCs exhibit bona fide LT-HSC activity

To investigate the developmental identity of *Hoxb5+* fetal liver (FL) HSCs, we utilized the previously described *Hoxb5*-mCherry reporter mouse, in which mCherry expression marks *Hoxb5*-expressing cells^24^. We first examined *Hoxb5*-mCherry expression across immunophenotypically defined hematopoietic populations at embryonic day E14.5. Within the FL, *Hoxb5*-mCherry expression was largely restricted to the HSC compartment, identified as Lineage⁻ c-Kit⁺ Sca1⁺ CD48⁻ CD150⁺ cells, and was largely absent from more differentiated progenitor populations (**Fig. 1A; Fig. S1A**). Confocal imaging of E14.5 FL sections further confirmed the presence of Hoxb5-mCherry⁺ c-Kit⁺ cells in situ (**Fig. 1B**). *Hoxb5+* HSCs were detected from E12.5 through E16.5, with peak frequency observed at E14.5 (**Fig. 1C**), indicating that *Hoxb5* expression could be maintained within a subset of fetal HSCs throughout the major period of liver hematopoietic expansion.

**Figure 1.**
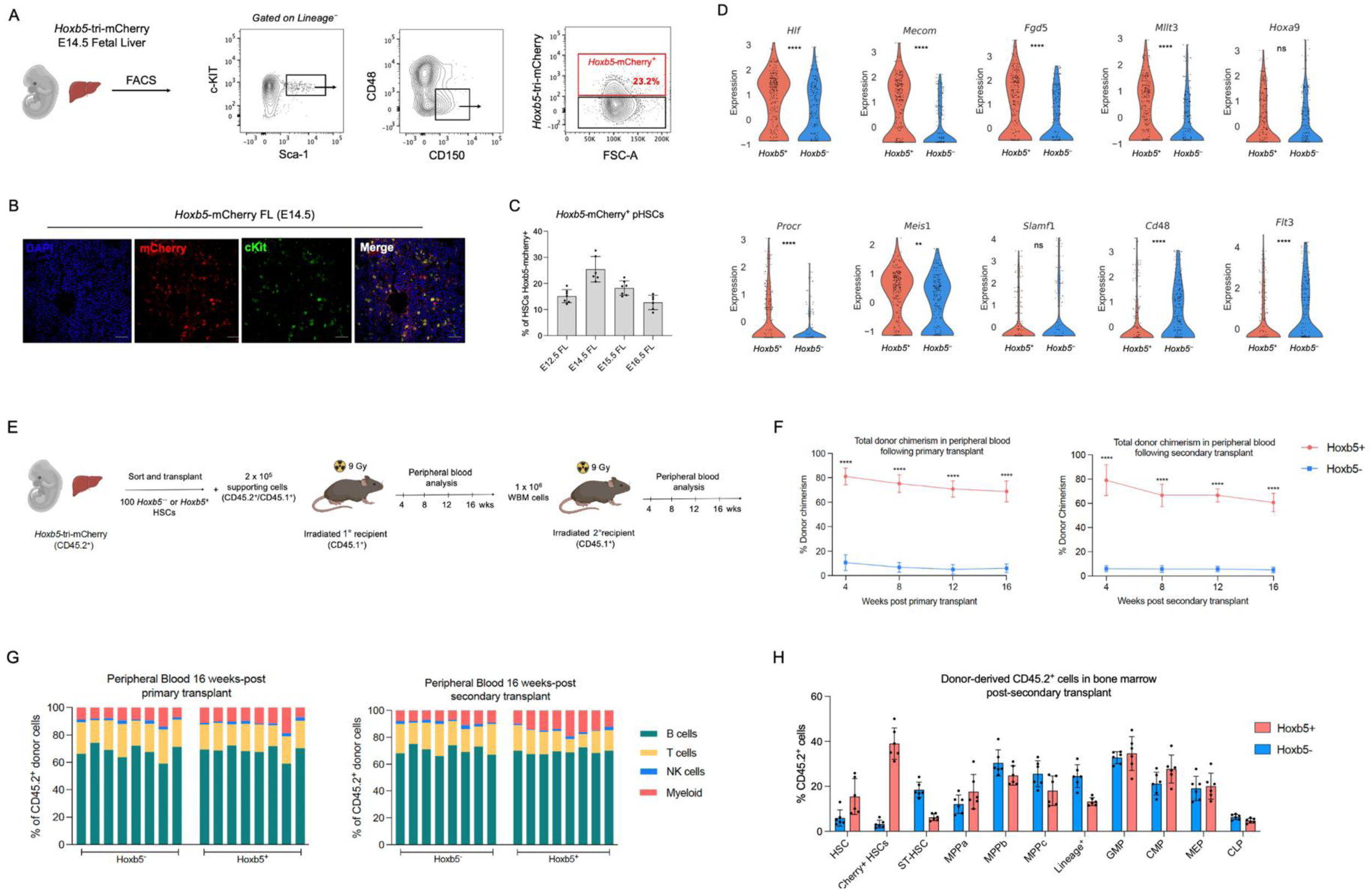
*Hoxb5+* FL HSCs exhibit bona fide LT-HSC activity. **(A)** Representative flow cytometry gating for identification of *Hoxb5*-mCherry⁺ HSCs in E14.5 fetal liver (FL) from *Hoxb5*-tri-mCherry embryos. **(B**) Representative confocal images of E14.5 FL sections from *Hoxb5*-tri-mCherry embryos showing DAPI, *Hoxb5*-mCherry and c-Kit staining. Scale bar = 50 μm. **(C)** Frequency of *Hoxb5*-mCherry⁺ cells among phenotypic HSCs in the FL across developmental stages (E12.5–E16.5). Each dot represents one embryo. Bars indicate mean ± SD. **(D)** Violin plots showing expression of representative LT-HSC-associated and progenitor-associated genes in sorted Hoxb5⁺ and Hoxb5⁻ FL HSCs by scRNA-seq. Each dot represents one cell. Statistical significance was determined using a two-sided Mann–Whitney U test. **(E)** Experimental schematic. One hundred *Hoxb5*⁺ or *Hoxb5*⁻ phenotypic HSCs were purified from E14.5 *Hoxb5*-tri-mCherry FLs (CD45.2) and transplanted together with Sca1⁻ radioprotective support cells into lethally irradiated adult recipients (CD45.1). **(F)** Total donor-derived peripheral blood chimerism following primary and secondary transplantation of *Hoxb5*⁺ or *Hoxb5*⁻ FL HSCs. Data are shown as mean ± SD (n = 5 recipients per group). **(G)** Peripheral blood lineage output at primary and secondary transplantation endpoints showing donor contribution to myeloid, B cell, T cell, and NK cell compartments. Each bar represents one recipient. **(H)** Donor-derived contribution to the bone marrow at the secondary transplantation endpoint. Each dot represents one recipient (n = 5). Bars indicate mean ± SD. ns, not significant; P values are denoted as ns, not significant; P < 0.05 (*), P < 0.01 (**), P < 0.001 (***), and P < 0.0001 (****).

We next asked whether *Hoxb5* expression distinguished a molecularly distinct FL HSC state. Single-cell RNA sequencing (scRNA-seq) of *Hoxb5*⁺ and *Hoxb5*⁻ FL HSCs revealed enrichment of canonical LT-HSC-associated genes, including *Hlf*, *Mecom*, *Fgd5*, *Mllt3*, *Procr*, and *Meis1*, within the Hoxb5⁺ population, whereas progenitor-associated genes such as *Cd48* and *Flt3* were preferentially expressed by Hoxb5⁻ HSCs (**Fig. 1D**). *Hoxb5*⁺ and *Hoxb5*⁻ FL HSCs exhibited comparable expression of fetal proliferative and developmental regulators, including *Mki67*, *Top2a*, *Lin28b*, and *Hmga2* (**Fig. S1B**), indicating that the transcriptional differences between the two populations were not simply attributable to differences in cell-cycle status or fetal identity. Both populations expressed hematopoietic markers such as Ptprc, whereas a greater fraction of Hoxb5⁺ HSCs expressed endothelial-associated genes including *Pecam1*, *Cdh5*, *Tek*, and *Kdr* (**Fig. S1C**). Together, these findings suggested that *Hoxb5*+ fetal HSCs are enriched for canonical LT-HSC transcriptional programs.

To determine whether *Hoxb5* expression also distinguished functionally definitive fetal LT-HSCs, we purified 100 *Hoxb5*⁺ or *Hoxb5*⁻ phenotypic HSCs from E14.5 *Hoxb5*-mCherry FLs and transplanted them into lethally irradiated adult recipients together with radioprotective support cells (**Fig. 1E**). Donor-derived peripheral blood chimerism was monitored for 16 weeks, after which whole bone marrow from primary recipients was transplanted into secondary irradiated recipients to assess long-term self-renewal capacity. *Hoxb5*⁺ FL HSCs generated robust hematopoietic reconstitution in both primary and secondary recipients, whereas *Hoxb5*⁻ HSCs contributed minimally throughout the duration of the experiment (**Fig. 1F, Fig. S1D-E**). Consistent with multilineage hematopoietic output, *Hoxb5*⁺ donor-derived grafts generated myeloid, B cell, T cell, and NK cell populations following both primary and secondary transplantation (**Fig. 1G**).

At the secondary transplantation endpoint, *Hoxb5*⁺ FL HSC-derived grafts exhibited substantially higher donor contribution within the bone marrow than *Hoxb5*⁻ grafts and reconstituted the phenotypic HSC compartment as well as downstream progenitor and mature hematopoietic populations, demonstrating hierarchical hematopoietic regeneration (**Fig. 1H**). Both *Hoxb5*-mCherry⁺ and Hoxb5-mCherry⁻ HSCs were detected among donor-derived HSCs, indicating that *Hoxb5* reporter expression is not uniformly maintained following transplantation. Together, these findings demonstrate that *Hoxb5*-expressing fetal liver HSCs possess robust serial multilineage repopulating activity, consistent with bona fide long-term hematopoietic stem cell function.

### E14.5 Hoxb5-expressing cells give rise to functional adult LT-HSCs

To determine whether *Hoxb5*-expressing embryonic cells contribute durably to adult hematopoiesis, we performed lineage tracing using *Hoxb5*-CreERT2;R26-ZsGreen mice. Pregnant dams were administered 4-hydroxytamoxifen (4-OHT) at E14.5, and labeled offspring were aged to 13 weeks prior to analysis (**Fig. 2A**).

**Figure 2.**
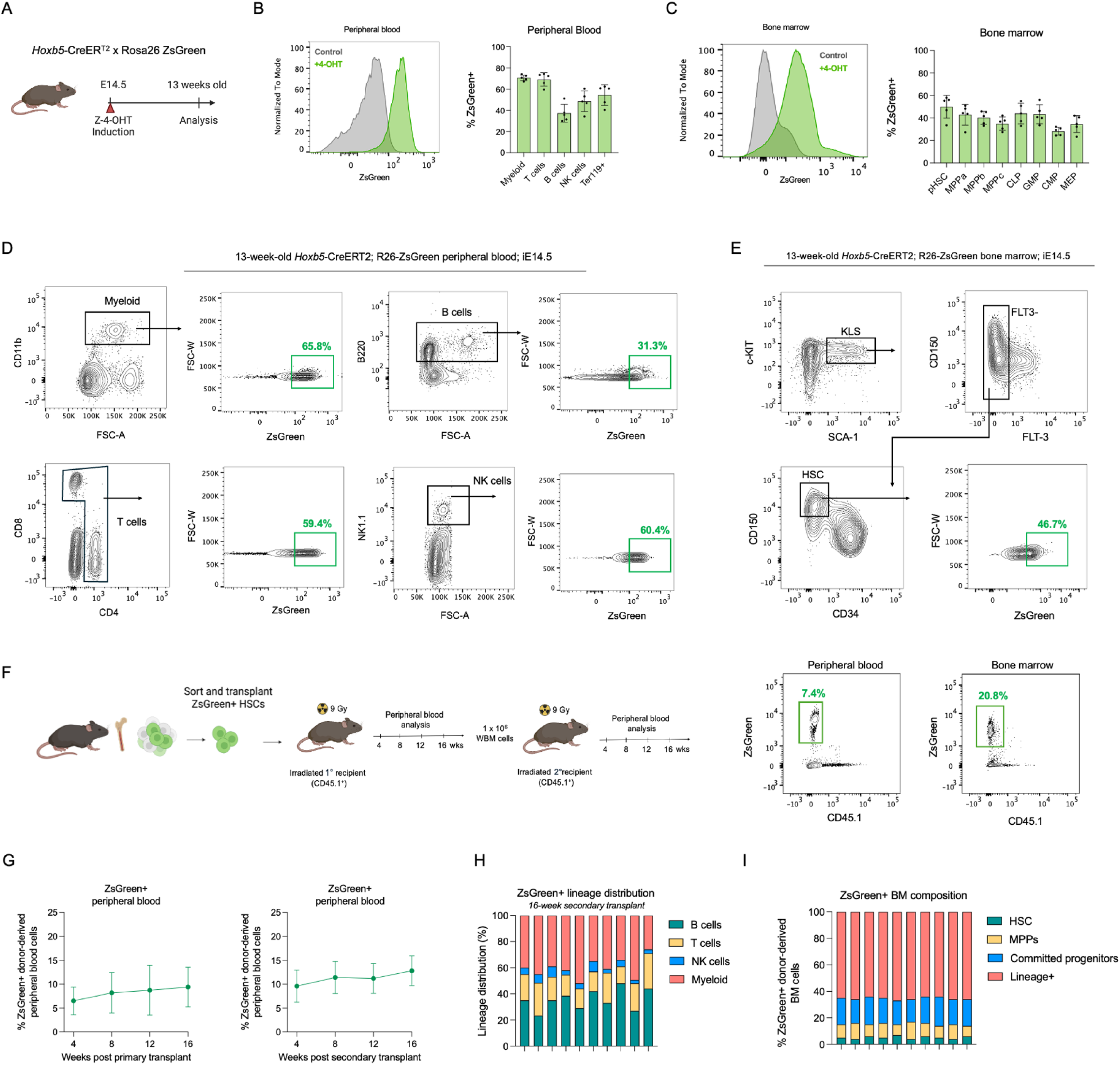
E14.5 *Hoxb5*-expressing cells contribute broadly to adult hematopoiesis and generate functional adult LT-HSCs. **(A)** Experimental schematic. Pregnant *Hoxb5*-CreERT2;R26-ZsGreen dams were administered 4-hydroxytamoxifen at *E14*.5 to label Hoxb5-expressing embryonic cells. Offspring were analyzed at 13 weeks of age. **(B)** Representative ZsGreen histograms of live peripheral blood cells from induced and non-induced control mice (left) and quantification of ZsGreen labeling within mature peripheral blood lineages (right), including myeloid cells, T cells, B cells, NK cells, and Ter119+ erythroid cells. **(C)** Representative ZsGreen histograms of live bone marrow cells from induced and non-induced control mice (left) and quantification of ZsGreen labeling across bone marrow hematopoietic stem and progenitor populations (right), including phenotypic HSCs (pHSCs), MPPa, MPPb, MPPc, CLP, GMP, CMP, and MEP populations. **(D)** Representative flow cytometric gating demonstrating ZsGreen labeling within adult peripheral blood lineages derived from E14.5 Hoxb5-expressing embryonic cells, including myeloid, B cell, T cell, and NK cell populations. **(E)** Representative flow cytometric gating demonstrating ZsGreen labeling within the adult bone marrow HSC compartment. Phenotypic HSCs were identified as Lineage− Sca1+ c-Kit+ FLT3− CD150+ CD34− cells and assessed for ZsGreen expression. **(F)** Experimental schematic of serial transplantation of ZsGreen+ phenotypic HSCs isolated from adult lineage-traced mice and representative flow plots showing ZsGreen+ cell chimerism in peripheral blood and bone marrow. **(G)** Analysis of ZsGreen+ donor-derived peripheral blood chimerism in primary and secondary recipients at 4, 8, 12, and 16 weeks post-transplant. Data are shown as mean ± SD (*n =* 10). **(H)** Lineage composition of donor-derived ZsGreen+ peripheral blood cells at 16 weeks following secondary transplantation and (**I)** distribution of donor-derived ZsGreen+ bone marrow cells across HSC, MPPs, progenitors (GMP, CMP, MEP, and CLP), and mature lineage compartments at the secondary transplant endpoint. Each bar represents an individual mouse.

ZsGreen labeling was detected in adult peripheral blood and bone marrow from induced animals but was absent from non-induced controls (**Fig. 2B-C**). In peripheral blood, E14.5 *Hoxb5*-expressing cells contributed to myeloid, erythroid, and lymphoid lineages, including CD11b+ myeloid cells, Ter119+ erythroid cells, CD3+ T cells, CD19+ B cells, and NK cells (**Fig. 2D**). Within the bone marrow, ZsGreen labeling was detected throughout the hematopoietic hierarchy, including phenotypic HSCs, MPP subsets, CLPs, GMPs, CMPs, and MEPs (**Fig. 2E**). These findings indicate that *Hoxb5*-expressing cells present at E14.5 contribute broadly to adult hematopoiesis and persist throughout the adult hematopoietic hierarchy, including the adult HSC compartment.

To determine whether adult HSCs derived from E14.5 *Hoxb5*-expressing cells retain long-term stem cell function, we purified ZsGreen+ phenotypic HSCs from adult lineage-traced mice and transplanted them into lethally irradiated recipients (**Fig. 2F**). ZsGreen+ HSCs generated donor-derived hematopoietic reconstitution in primary recipients and maintained engraftment following secondary transplantation (**Fig. 2G**). Donor-derived cells exhibited multilineage output in peripheral blood (**Fig. 2H**) and reconstituted HSC, progenitor, and mature hematopoietic compartments in the bone marrow at the secondary transplant endpoint (**Fig. 2I**). Together, these findings demonstrate that *Hoxb5*-expressing cells present at E14.5 give rise to adult LT-HSCs that retain serial multilineage repopulating activity, establishing a direct developmental link between embryonic *Hoxb5*-expressing HSCs and the adult long-term stem cell compartment.

### Fetal *Hoxb5*+ HSCs maintain LT-HSC function despite extensive proliferative activity

To investigate how *Hoxb5*⁺ HSCs mature across development, we performed scRNA-seq analysis of *Hoxb5*⁺ HSCs isolated from FL at E12.5, E14.5, E15.5, and E16.5 together with adult BM *Hoxb5*⁺ HSCs (**Fig. 3A, Fig. S2A**). Integrated analysis revealed a clear developmental separation between fetal liver and adult bone marrow *Hoxb5*⁺ HSCs, indicating substantial remodeling of HSC state during maturation.

**Figure 3.**
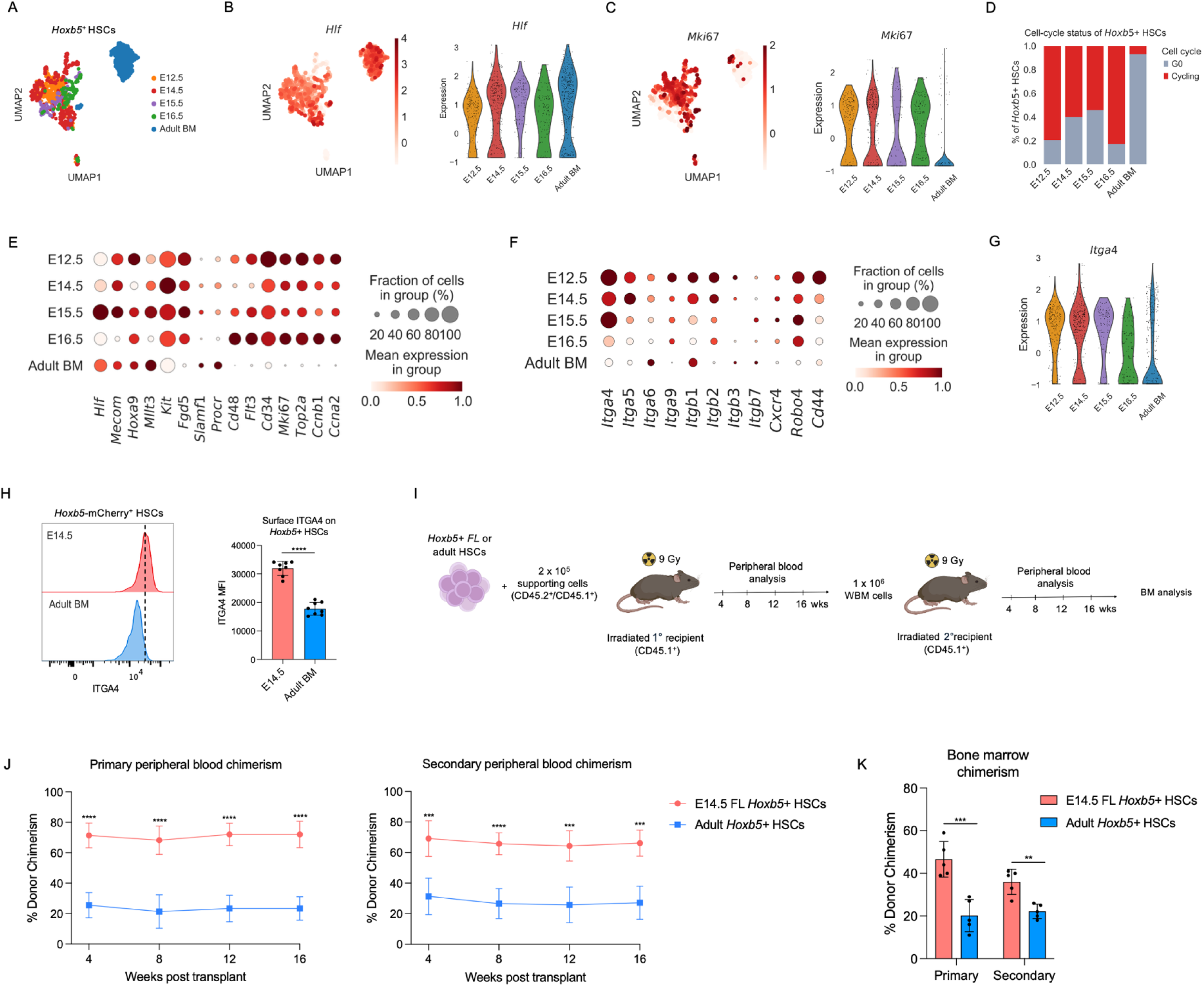
**Cycling fetal *Hoxb5***⁺ **HSCs retain robust long-term self-renewal and engraftment potential (A)** scRNA-seq UMAP of *Hoxb5*+ HSCs sorted from E12.5, E14.5, E15.5, and E16.5 FL and 6-week-old adult bone marrow (BM). **(B-C)** UMAP visualization and violin plot showing expression of the *Hlf* and *Mki67* across developmental stages. **(D)** Cell-cycle status of *Hoxb5*+ HSCs across development inferred from scRNA-seq data. Proportions of cells in cycling and G0 states are shown for each developmental stage. **(E)** Dot plot showing relative expression of canonical HSC-associated genes and cell-cycle-associated genes across developmental stages in Hoxb5⁺ HSCs. Dot size indicates the fraction of expressing cells and color intensity indicates mean expression. **(G)** Violin plot showing Itga4 expression across developmental stages in *Hoxb5*⁺ HSCs. **(H)** Representative flow cytometry histograms and quantification of surface ITGA4 expression on E14.5 FL and adult BM Hoxb5⁺ HSCs. Dashed line indicates peak ITGA4 staining intensity. Each dot represents one embryo or mouse. Quantification is shown as mean fluorescence intensity (MFI). **(I)** Experimental schematic for serial transplantation of sorted E14.5 FL or adult BM Hoxb5⁺ HSCs into lethally irradiated primary recipients followed by secondary transplantation. **(J)** Peripheral blood donor chimerism in primary and secondary transplant recipients over time following transplantation of fetal liver or adult bone marrow *Hoxb5*⁺ HSCs. **(K)** Bone marrow donor chimerism at endpoint analysis in primary and secondary recipients transplanted with fetal liver or adult bone marrow *Hoxb5*⁺ HSCs. Data are shown as mean ± SEM. Statistical significance was determined using Mann–Whitney U tests unless otherwise indicated. P values are denoted as ns, not significant; P < 0.05 (*), P < 0.01 (**), P < 0.001 (***), and P < 0.0001 (****).

Consistent with a hematopoietic program, both fetal and adult *Hoxb5*⁺ HSCs expressed high levels of *Hlf*, *Mecom*, *Hoxa9, and Mllt3* (**Fig. 3B, Fig. S2A-B**). Fetal Hoxb5⁺ HSCs exhibited widespread expression of the proliferation-associated gene Mki67, whereas the majority of adult BM Hoxb5⁺ HSCs lacked detectable Mki67 expression (**Fig. 3C, Fig S2C**). Cell-cycle inference further demonstrated that the majority of fetal Hoxb5⁺ HSCs occupied actively cycling states, whereas adult *Hoxb5*⁺ HSCs were predominantly quiescent **(Fig. 3D**). Despite their extensive proliferative activity, fetal Hoxb5⁺ HSCs maintained expression of canonical LT-HSC-associated genes including *Hlf*, *Mecom*, *Hoxa9*, *Mllt3*, *Kit*, *Fgd5*, *Slamf1*, and *Procr* across developmental stages (**Fig. 3E**). These findings indicate that fetal *Hoxb5*⁺ HSCs maintain an LT-HSC transcriptional program while undergoing extensive proliferative expansion.

Given the established roles of adhesion molecules in HSC migration, homing, and niche interactions, as well as prior evidence that fetal and adult hematopoietic stem and progenitor cells exhibit distinct adhesive and migratory properties^25–29^, we next examined integrin and adhesion-associate gene expression across development. Multiple adhesion-associated genes were enriched in fetal *Hoxb5*⁺ HSCs, with *Itga4* remaining highly expressed throughout fetal stages relative to adult BM *Hoxb5*⁺ HSCs (**Fig. 3F-G, Fig. S2D**). Consistent with these transcriptional findings, E14.5 FL *Hoxb5*⁺ HSCs expressed substantially higher levels of surface ITGA4 protein than adult BM *Hoxb5*⁺ HSCs (**Fig.34H**).

To determine whether extensive proliferation was associated with reduced long-term stem cell function, we directly compared the regenerative activity of fetal and adult *Hoxb5*⁺ HSCs by serial transplantation. Equal numbers of E14.5 FL or adult BM *Hoxb5*⁺ HSCs were transplanted into lethally irradiated recipients and followed through serial transplantation (**Fig. 3I**). Despite their highly proliferative transcriptional state, fetal liver *Hoxb5*⁺ HSCs generated substantially higher donor chimerism than adult bone marrow *Hoxb5*⁺ HSCs in both primary and secondary recipients (**Fig. 3J**). At endpoint analysis, fetal liver *Hoxb5*⁺ HSC-derived grafts also produced significantly greater bone marrow donor chimerism than adult *Hoxb5*⁺ HSC-derived grafts (**Fig. 3K**). To determine whether this developmental program is conserved in humans, we analyzed Smart-seq3 profiles of human fetal liver HSCs (11 gestational weeks) and adult bone marrow HSCs. Human fetal liver HSCs expressed canonical HSC regulators including *HLF*, *MECOM*, *HOXA9*, *MLLT3*, and *SPINK2*, and exhibited elevated *MKI67* and *ITGA4* expression relative to adult bone marrow HSCs (**Fig. S3A-C**), consistent with conservation of a proliferative, adhesion-enriched fetal HSC state across species.

Together, these findings demonstrate that fetal *Hoxb5*⁺ HSCs retain robust long-term self-renewal and regenerative capacity despite extensive proliferative activity, indicating that acquisition of LT-HSC function precedes establishment of the quiescent adult HSC state.

### ITGA4-mediated adhesion contributes to fetal HSC engraftment

Given the elevated expression of ITGA4 in fetal *Hoxb5*⁺ HSCs, we next sought to identify candidate adhesion partners within the developing fetal liver microenvironment. We performed scRNA-seq on sorted HSPCs together with CD45⁻ fetal liver cells enriched for endothelial and stromal populations based on cell surface expression of CD31, CD144, TIE2, VEGFR2, or PDGFRA (**Fig. S4A).** Unsupervised clustering identified endothelial, stromal, EMP-like, and HSPC populations within the developing fetal liver niche (**Fig. 4A**, **Fig S4B**). Among these populations, *Vcam1* expression was enriched within endothelial cells (**Fig. 4B, Fig. S4C**). Endothelial *Vcam1* expression was highest during mid-gestation and progressively declined at later developmental stages (**Fig. 4C**). Consistent with these transcriptional findings, flow cytometry analysis demonstrated that a substantial fraction of CD45⁻CD144⁺ endothelial cells expressed VCAM1 between E12.5 and E15.5, followed by a reduction at E16.5-E18.5 (**Fig. 4D**). These findings identify fetal liver endothelial cells as a prominent source of VCAM1 during the developmental window of active fetal HSC expansion.

**Figure 4.**
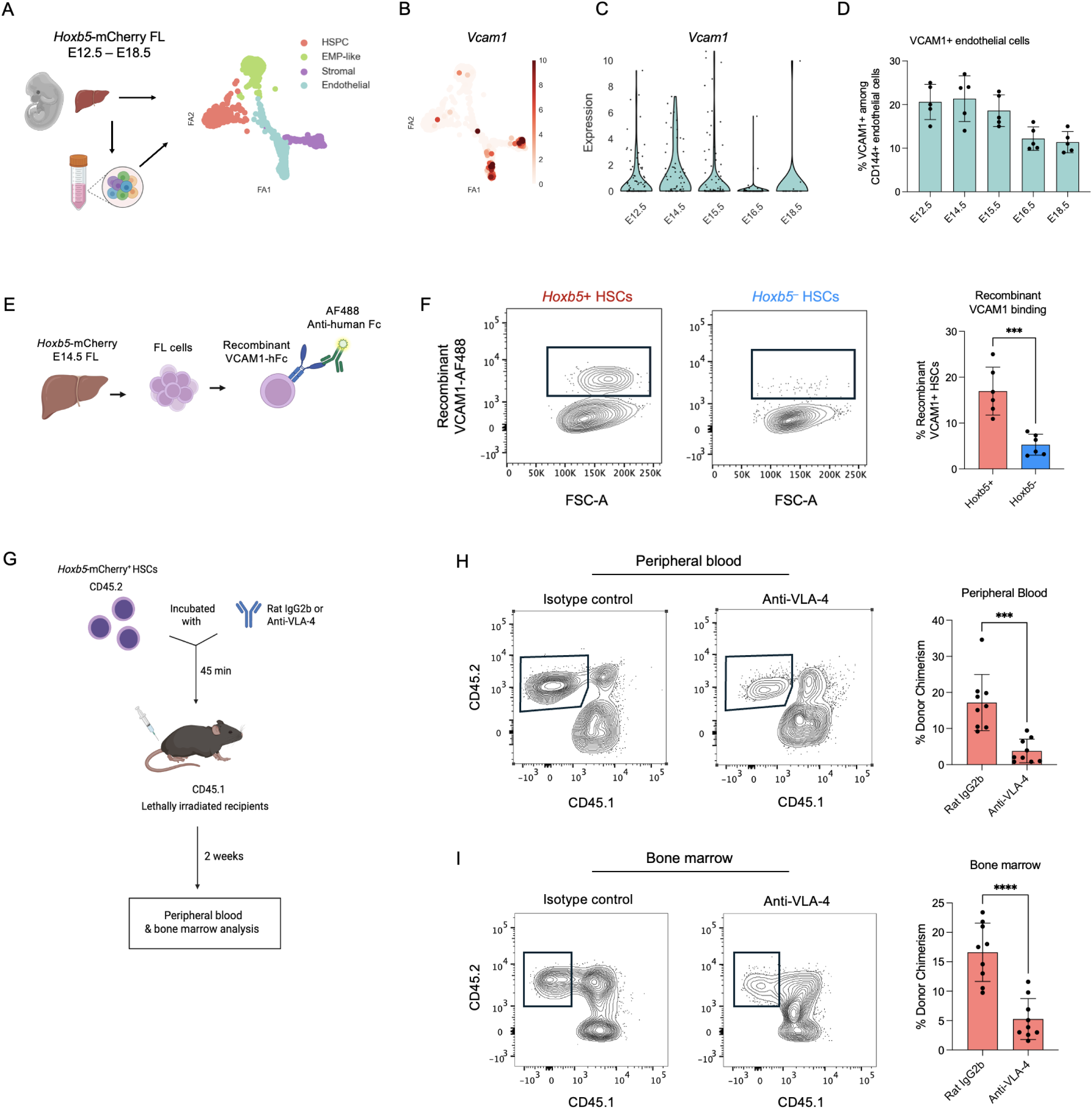
ITGA4-mediated adhesion supports fetal HSC engraftment. **(A)** Experimental schematic and integrated scRNA-seq analysis of HSPCs and CD45^-^ fetal liver cells isolated from E12.5-E18.5 *Hoxb5*-tri-mCherry embryos. UMAP visualization identifies endothelial, stromal, EMP-like, and HSPC populations within the developing fetal liver microenvironment. **(B)** UMAP visualization showing *Vcam1* expression within fetal liver niche populations. **(C)** Violin plot showing *Vcam1* expression within endothelial cells across developmental stages. **(D)** Flow cytometry quantification of VCAM1+ cells among CD45^-^ CD144+ endothelial cells across fetal liver developmental stages. **(E)** Experimental schematic for recombinant VCAM1 binding assay. Single-cell suspensions prepared from E14.5 *Hoxb5*-mCherry FLs were incubated *ex vivo* with recombinant VCAM1-hFc followed by AF488-conjugated anti-human Fc secondary antibody staining. **(F)** Representative flow cytometry plots and quantification of recombinant VCAM1 binding by *Hoxb5*+ and *Hoxb5*^-^ fetal liver HSCs. **(G)** Experimental schematic for *ex vivo* VLA-4 blockade prior to transplantation. Purified E14.5 Hoxb5-mCherry+ FL HSCs were incubated with anti-VLA-4 or Rat IgG2b isotype control antibodies prior to transplantation into lethally irradiated recipients. **(H)** Representative peripheral blood flow cytometry plots and quantification of donor chimerism two weeks after transplantation of anti-VLA-4-or isotype-treated fetal *Hoxb5*+ HSCs. **(I)** Representative bone marrow flow cytometry plots and quantification of donor chimerism within the bone marrow HSC compartment 4 weeks after transplantation of anti-VLA-4-or isotype-treated fetal *Hoxb5*+ HSCs. Data are shown as mean ± SEM. Statistical significance was determined using Mann–Whitney U tests. *P* values are denoted as ns, not significant; *P* < 0.05 (\**), P < 0.01 (**), P < 0.001 (\*\*\**), and *P* < 0.0001 (****).

We next asked whether *Hoxb5*⁺ fetal HSCs exhibit enhanced VCAM1-binding capacity. Single-cell suspensions prepared from E14.5 *Hoxb5*-mCherry fetal livers were incubated *ex vivo* with recombinant VCAM1-hFc, and binding was detected using a fluorescent anti-human Fc secondary antibody (**Fig. 4E**). *Hoxb5*⁺ fetal HSCs displayed substantially greater recombinant VCAM1 binding than *Hoxb5*⁻ fetal HSCs (**Fig. 4F**).

To determine whether ITGA4-mediated adhesive interactions contribute functionally to fetal HSC engraftment, purified E14.5 *Hoxb5*⁺ fetal liver HSCs were incubated *ex vivo* with blocking anti-VLA-4 antibodies or isotype control antibodies prior to transplantation into lethally irradiated recipients (**Fig. 4G**). VLA-4 blockade markedly reduced donor-derived peripheral blood chimerism relative to isotype-treated controls 4 weeks after transplantation (**Fig. 4H**). Consistent with impaired engraftment, donor contribution to the bone marrow HSC compartment was also significantly reduced following VLA-4 blockade (**Fig. 4I**). Together, these findings identify the ITGA4–VCAM1 adhesion axis as an important regulator of fetal *Hoxb5*⁺ HSC engraftment and establish a functional role for ITGA4-mediated adhesion during fetal hematopoiesis.

## Discussion

HSCs undergo major developmental transitions as they mature from rapidly expanding fetal populations into predominantly quiescent adult stem cells^12–15^. While fetal and adult HSCs have long been recognized to differ in proliferative activity^20–23^, the developmental relationship between these populations and the mechanisms that support fetal HSC function remain incompletely understood. In this study, we characterize a population of *Hoxb5*-expressing fetal liver HSCs that exhibit multilineage and serial repopulating activity and demonstrate that these cells contribute directly to the adult LT-HSC compartment. Our findings further reveal that acquisition of long-term stem cell function precedes establishment of the quiescent adult HSC state and identify enhanced ITGA4-mediated adhesion as a prominent feature of fetal LT-HSCs.

Prior studies established Hoxb5 as a highly selective marker of adult LT-HSCs and demonstrated that Hoxb5-expressing LT-HSCs make major contributions to steady-state hematopoiesis in vivo^24,30^. However, whether *Hoxb5*-expressing cells contribute to a developmentally conserved LT-HSC population spanning fetal and adult hematopoiesis remained unclear. Here, we show that *Hoxb5*-expressing fetal liver HSCs are enriched for canonical LT-HSC transcriptional programs and possess multilineage and serial repopulating activity. Importantly, embryonic lineage tracing demonstrated that *Hoxb5*-expressing cells present during fetal liver hematopoiesis contribute to the adult HSC compartment, and that adult HSCs derived from these cells retain serial multilineage repopulating activity. Together, these findings establish a direct developmental relationship between fetal *Hoxb5*-expressing HSCs and adult LT-HSCs and suggest that key features of long-term stem cell identity are established during fetal life.

These findings have important implications for our understanding of HSC ontogeny. The fetal liver is generally viewed as a transient site of HSC expansion that seeds the bone marrow shortly before birth^8,12^. The fetal liver is generally viewed as a transient site of HSC expansion that supports the rapid amplification of the stem cell pool before bone marrow colonization. Our results demonstrate that *Hoxb5*⁺ LT-HSCs present during this developmental stage give rise to adult LT-HSCs with self-renewal and regenerative capacity. Thus, at least a subset of fetal liver HSCs are not merely transient developmental intermediates but instead contribute directly to the long-lived stem cell compartment that sustains lifelong hematopoiesis.

A central observation of our study is that fetal LT-HSCs maintain exceptional regenerative potential despite extensive proliferative activity. Previous studies have established that fetal liver HSCs undergo rapid expansion during development, whereas adult HSCs are predominantly maintained in a quiescent state^21–23,31–33^. Consistent with these observations, fetal *Hoxb5*⁺ HSCs exhibited widespread cell-cycle activity, while adult *Hoxb5*⁺ HSCs were largely quiescent. Despite these marked differences in proliferative status, fetal *Hoxb5*⁺ HSCs consistently outperformed adult *Hoxb5*⁺ HSCs in both primary and secondary transplantation assays. These findings suggest that the relationship between proliferation and stem cell fitness is developmentally regulated. Rather than being incompatible with long-term stem cell function, proliferation during fetal life appears to be integrated into a physiological developmental program that expands the stem cell pool while preserving self-renewal and regenerative capacity. More broadly, our findings suggest that acquisition of long-term stem cell function precedes establishment of the quiescent adult HSC state.

Our data further identify the ITGA4–VCAM1 adhesion axis as a prominent feature of this fetal HSC state. VLA-4/VCAM1 and related integrin pathways have long been implicated in hematopoietic stem and progenitor cell adhesion, homing, lodgment, and retention^29,31,32^. Prior studies demonstrating roles for α4 and β1 integrins during fetal and adult hematopoiesis^25,28,34–39^. In this context, our findings extend this framework to *Hoxb5*-expressing fetal LT-HSCs by showing that these cells maintain elevated ITGA4 expression, preferentially bind recombinant VCAM1, and require VLA-4 activity for efficient engraftment. In parallel, fetal liver endothelial cells expressed high levels of VCAM1 during the developmental window in which fetal Hoxb5⁺ HSCs were most abundant. Analysis of human fetal liver HSCs revealed elevated *ITGA4* expression relative to adult bone marrow HSCs, suggesting that aspects of the fetal adhesion program identified here may be conserved across species. Together, these findings suggest that ITGA4-mediated adhesive interactions are a defining feature of fetal *Hoxb5*⁺ HSCs and may contribute to their enhanced engraftment competence during developmental hematopoiesis.

Several important questions remain unresolved. Although our transplantation studies demonstrate a requirement for VLA-4 during engraftment, whether ITGA4 signaling contributes directly to endogenous fetal liver HSC maintenance remains to be determined. Recent studies have identified specialized endothelial, stromal, and macrophage populations that support developmental hematopoiesis, raising the possibility that multiple niche-derived signals cooperate to maintain fetal HSC self-renewal during periods of rapid expansion^40–45^. Defining how fetal HSCs preserve long-term self-renewal while actively cycling may provide insight into developmental programs that uncouple proliferation from loss of hematopoietic stem cell function and could potentially be leveraged to enhance adult HSC expansion and regenerative capacity.

The ability of fetal HSCs to maintain robust regenerative capacity despite extensive proliferation stands in contrast to the functional decline often associated with HSC cycling during aging, inflammation, and hematopoietic stress^46–49^. Understanding how fetal HSCs uncouple proliferation from loss of stem cell function may therefore provide insights into strategies for HSC expansion, transplantation, and rejuvenation. It is also notable that several hematopoietic populations with limited engraftment potential, including pre-circulation embryonic HSC precursors and human pluripotent stem cell-derived phenotypic HSCs, fail to robustly reconstitute adult recipients despite expressing many features associated with definitive HSC identity^50–52^. Our findings raise the possibility that acquisition of adhesive programs involving ITGA4 may represent an additional developmental step associated with functional engraftment competence during HSC maturation.

In summary, our findings demonstrate that *Hoxb5*-expressing fetal liver HSCs contribute directly to the adult LT-HSC compartment and reveal a developmental continuum linking fetal and adult long-term hematopoietic stem cells. These studies further show that LT-HSC function is established prior to acquisition of adult quiescence and identify enhanced ITGA4-mediated adhesion as a characteristic feature of the fetal stem cell state. Together, these findings provide new insight into the developmental programs that generate and maintain lifelong hematopoiesis.

## Supplementary Figure Legends

**Supplementary Figure 1. Characterization of *Hoxb5*-mCherry expression and transcriptional features of fetal liver HSCs. (A)** Representative flow cytometry analysis of *Hoxb5*-mCherry expression across immunophenotypically defined hematopoietic populations in E14.5 fetal liver. *Hoxb5*-mCherry expression was enriched within the Lineage⁻ c-Kit⁺ Sca1⁺ CD48⁻ CD150⁺ HSC compartment and largely absent from Lineage⁺ cells, Lineage⁻ c-Kit⁺ Sca1⁻ progenitors, KLS CD48⁻ CD150⁻ multipotent progenitors, and KLS CD48⁺ CD150⁻ progenitors. Histograms show Hoxb5-mCherry fluorescence intensity within the indicated populations. FL cells from C57/BL6 mice with no mCherry reporter are shown for comparison. **(B)** Violin plots showing scRNA-seq expression of proliferation-and fetal-associated genes (*Mki67*, *Top2a*, *Lin28b*, and *Hmga2*) in E14.5 fetal liver *Hoxb5*⁺ and *Hoxb5*⁻ HSCs. **(C)** Expression of hematopoietic and endothelial-associated genes in E14.5 fetal liver *Hoxb5*⁺ and *Hoxb5*⁻ HSCs. *Ptprc* (CD45), *Cdh5* (VE-cadherin), *Pecam1* (CD31), *Tek* (Tie2) and *Kdr* (VEGFR2). Statistical significance was determined using a two-sided Mann–Whitney U test. ns = not significant; P < 0.05 (), P < 0.01 (), P < 0.001 (), and P < 0.0001 (****).

**Supplementary Figure 2. Developmental transcriptional profiling of fetal liver and adult *Hoxb5***⁺ **HSCs. (A)** UMAP visualization of Hoxb5⁺ HSCs and feature plots showing expression of representative HSC-associated genes. **(B)** Dot plot showing expression of LT-HSC-associated genes and proliferation-associated genes across developmental stages. **(C)** Cell-cycle phase assignment (G0, G2M, S) of *Hoxb5*⁺ HSCs across development determined by scRNA-seq. **(D)** Violin plots showing *Itga4* expression in cycling (*Mki67*⁺) and non-cycling (*Mki67*⁻) *Hoxb5*⁺ HSCs at each developmental stage. Statistical significance was determined using a two-sided Mann–Whitney U test. P < 0.0001 (****).

**Supplementary Figure 3. Human fetal liver HSCs exhibit conserved fetal stem cell and adhesion programs. (A)** UMAP visualization of Smart-seq3 profiles from immunophenotypically defined human HSCs isolated from 11 gestational week (GW) fetal liver (FL) and adult bone marrow (BM). Feature plots show expression of canonical HSC-associated genes (HLF, *MECOM*, *HOXA9*, *RUNX1*, *MLLT3*, *SPINK2*, and *PROM1*), as well as *HOXB5* and the proliferation-associated gene *MKI67*. **(B)** Dot plot showing expression of adhesion-and homing-associated genes in human fetal liver and adult bone marrow HSCs. **(C)** Violin plot showing *ITGA4* expression in human fetal liver and adult bone marrow HSCs. Statistical significance was determined using a two-sided Mann–Whitney U test, ****P < 0.0001.

**Supplementary Figure 4. Identification of fetal liver niche populations and sorting strategy for niche scRNA-seq analysis. (A)** UMAP representation of single-cell RNA sequencing data generated from fetal liver hematopoietic stem and progenitor cells (HSPCs) together with CD45⁻ fetal liver cells enriched for endothelial and stromal populations by flow cytometric sorting. Unsupervised clustering identified HSPC, endothelial, stromal, and EMP-like populations. Dot plot shows expression of representative marker genes used for cluster annotation. **(B)** Feature plots showing expression of *Hlf*, *Kit*, *Itga2b*, *Cdh5*, *Pecam1*, *Tek*, *Pdgfra*, and *Vcam1* across the integrated fetal liver dataset. **(C)** Representative flow cytometry plots showing the gating strategy used to isolate fetal liver niche populations for scRNA-seq analysis. CD45⁻ fetal liver cells expressing endothelial-associated markers (CD31, CD144, TIE2, VEGFR2, and VCAM1) or the stromal marker PDGFRA were collected and subjected to scRNA-seq.

## Methods

### Mice

*Hoxb5*–tri-mCherry (CD45.2 C57BL/6J background) mice were used as donor cells for transplantation as well as for analysis. *Hoxb5*-CreERT2 mice were crossed with R26-ZsGreen mice and offspring were used for lineage tracing experiments. Mice were bred at our animal facility according to NIH guidelines. 8-to-12-week old female B6.SJL-Ptprca Pepcb/BoyJ mice (Jackson Laboratory) were used as recipients for transplantation assays. Supporting cells were collected from B6.SJL-Ptprca Pepcb/BoyJ□×□C57BL/6J (CD45.1+/CD45.2+). All animal protocols were approved by the Stanford University Administrative Panel on Laboratory Animal Care.

### Isolation of fetal liver and bone marrow cells

Fetal livers were carefully dissected from embryos using fine forceps under a dissection microscope to ensure precision and minimize tissue damage. The dissected liver tissues were then finely minced and incubated at 37C for 30min in pre-warmed digestion media containing M199 media (Thermo), 0.1% Pluronic’s 188 (Thermo), 2mg/mL Colleganse II (Thermo), 0.2mg/mL DNAseI, and 0.1% PVA (Thermo). Liver tissues were then gently passed through a 100 μm filter to remove any debris and clumps of tissue, ensuring a single-cell suspension. Following filtration, the cells were washed twice in PBS with 1% PVA (FACS buffer) to remove any residual contaminants and prepare them for antibody staining.

For bone marrow isolation, the tibia, femur, and pelvis were carefully dissected from the mice. The bones were then cleaned and crushed using a mortar and pestle in FACS buffer. After crushing, the supernatant containing the cellular components was collected for further processing. Red blood cells were depleted by incubating cells with ACK lysis buffer (Thermo) for 10 mins at room temperature, followed by centrifugation to pellet the remaining cells. Cells were Fc-blocked with TruStain FcX (BioLegend) for 10 minutes prior to staining with antibodies. To isolate c-KIT+ cells, the samples were further incubated with anti-c-KIT APC-eFluor780 (Thermo) for 20 mins at 4C. Following this incubation, the cells were washed twice with FACS buffer. After washing, the cells were incubated with anti-APC beads for 10 mins at 4C to enable the separation of c-KIT+ cells. The c-KIT-enriched cells were then isolated using LS magnetic columns (Miltenyi Biotec) according to the manufacturer’s protocol.

To isolate supporting cells from CD45.1+CD45.2+ mice, bone marrow was first harvested as previously described. Following isolation, bone marrow cells were incubated with SCA depletion beads (Miltenyi Biotec). The SCA depletion beads were added in accordance with the manufacturer’s instructions. After incubation, the cell suspension was washed and transferred to LS magnetic columns (Miltenyi Biotec) to facilitate the separation of SCA-depleted cells. The flow-through fraction, containing the Sca1-depleted supporting cells, was collected for transplantation.

### Isolation of human fetal liver and adult bone marrow cells

Human fetal liver (11 gestational weeks) was obtained Advanced Bioscience Resources (ABR, Alameda, CA) and shipped overnight in BioWhittaker RPMI-1640 media supplemented with L-glutamine.. Fetal liver tissue was mechanically dissociated using sterile razor blades and incubated in M199 medium supplemented with 0.2mg/mL DNase I and 2.2mg/mL Collagenase Type II, 0.1% Pluronics, and 0.1% BSA to generate a single-cell suspension. Following enzymatic digestion, cells were filtered through a 70-μm cell strainer, washed with FACS buffer (PBS containing 2% fetal bovine serum and 2 mM EDTA), and stained with fluorophore-conjugated antibodies.

Adult human bone marrow samples were obtained from femoral head and neck specimens collected during elective total hip arthroplasty procedures under protocols approved by the appropriate Institutional Review Board. Bone marrow cells were harvested by mechanically scraping the bone surfaces of the femoral neck and proximal femur, followed by suspension in cold staining buffer. Cell suspensions were filtered through a 70-µm cell strainer to remove debris and bone fragments, washed with FACS buffer, and stained with fluorophore-conjugated antibodies for flow cytometric analysis and sorting.

### Flow cytometry and cell sorting

Flow cytometry and cell sorting were performed on a FACS Aria II cell sorter (BD Biosciences) and analyzed using FlowJo software. As described previously, we defined HSCs in the fetal liver as: Lineage-Sca1+ Kit+ (KLS) CD150+ CD48-. HSCs in the adult bone marrow defined as: Lineage-Sca1+ Kit+ (KLS) CD150+ FLT3-CD34-. The following antibodies were included in the HSC and progenitor panel: c-Kit (APC-eFluor870), Sca-1 (BV395), CD150 (BV640), CD48 (AF700), CD16/32 (BV711), IL7RA (BV510), CD34 (FITC), and ITGA4 (PE). Exclusion using the following cocktail of biotinylated “lineage” antibodies were also used to identify HSCs: Ter119, CD4, CD8, B220, Gr1 and NK1.1, followed by secondary staining with BV737-conjugated Streptavidin. The following antibodies were included to capture non-hematopoietic cells in the FL: CD45 (BV737) for immune cell exclusion, CD31 (APC-Cy7), CD144 (PE-Cy7), TIE2 (BUV395), PDGFRA (BV605), and VCAM1 (PerCP-Cy5.5). Samples were incubated with anti-mouse CD16/32 Fc Block (BD Biosciences) for 10 minutes prior to staining with antibodies. Antibody staining was performed at 4 °C and cells were incubated for 30 min. Before sorting or analysis, cells were stained with SYTOX Red Dead Cell Stain (Life Technologies) to assess viability as per the manufacturer’s recommendations.

The following antibodies were included in the human HSC and progenitor panel: CD34 (APC-Cy7), CD38 (BV650), CD45RA (BV785), CD90 (FITC), CD11b (BV711), IL7RA (AF700), FLT3 (PerCP-Cy5.5). Exclusion using the following cocktail of “lineage” antibodies in PE-Cy5 were also used to identify HSPCs: CD3, CD4, CD8. CD14, CD19, CD20, CD56, and CD235a. Samples were incubated with Human TruStain FcX (Fc Receptor Blocking Solution) for 10 minutes prior to staining with antibodies. Antibody staining was performed at 4 °C and cells were incubated for 30 min. Before sorting or analysis, cells were stained with Proprium Iodide to assess viability as per the manufacturer’s recommendations.

### HSC transplants and peripheral blood analysis

B6.SJL-Ptprc Pepcb/BoyJ (Jackson Laboratory) recipient mice received a split dose of irradiation (two doses of 4.5Gy, separated by 4 hours) for a total dosage of 9Gy 24h before transplantation. For transplantation assays, 100 sorted donor *Hoxb5*+ or *Hoxb5*⁻ HSCs (KLS CD150+ CD48-Hoxb5-tri-mCherry+/-) from Hoxb5-tri-mCherry mice were first combined with 2×105 Sca-1 depleted whole bone marrow cells from B6.SJL-Ptprca Pepcb/BoyJ□×□C57BL/6J F1 mice (CD45.1+/CD45.2+) in 200 μl of PBS with 2% FBS, then injected into the retro-orbital venous plexus. Peripheral blood analyses were performed at 4, 8, 12, and 16 weeks after primary and secondary transplantations. At each time point, 50uL of blood was collected retro-orbitally and added to 500uL PBS with 2mM EDTA. Red blood cells were lysed using ACK lysing buffer (Thermo) for 5 min at room temperature followed by blocking with anti-mouse CD16/32 mouse BD Fc Block (BD Biosciences). Leukocytes were stained with antibodies against CD45.2 (BV421), CD45.1 (BV785), Ter119 (BV737), CD11b (FITC), Gr-1 (BV395), CD8 (PE), CD4 (PE-Cy7), NK1.1 (PE-Cy5), and B220 (APC-Cy7). For each mouse, the percentage of donor chimerism in the peripheral blood was defined as the percentage of CD45.1-CD45.2+ cells among total cells.

At the end of secondary transplantation, mice were euthanized and bone marrow from femurs, tibias, and pelvises were prepared as described above. For HSC and progenitor analysis, cells were stained with the following antibodies: CD45.2 (BV421), CD45.1 (BV785), c-KIT (APC-eFluor870), Sca-1 (BV395), CD150 (BV650), FLT3 (PerCP-eFluor710), CD41 (BV510), CD34 (BV711), CD16/32 (PE), and IL7Ra (AF700). A lineage cocktail comprising biotin-conjugated antibodies targeting Ter119, CD4, CD8, Gr-1, B220, and NK1.1 was used to exclude differentiated cells. Lineage-positive cells were subsequently stained with BV737-conjugated Streptavidin.

### Tissue staining and confocal imaging

*Hoxb5*-tri-mCherry embryos were dissected at embryonic day E14.5. Fetal livers were embedded in O.C.T. for sectioning and stored at-80C. Before sectioning, fetal livers were fixed in 4% P.F.A. for 35 minutes and washed 3x in PBS with 30 minutes in between each wash. For immunohistochemistry, 7 um fetal livers are placed in 5% donkey serum with 0.5% tritonX and PBS mix for 2 hours at room temperature. Primary antibodies were added to a 1% B.S.A. with 0.5% tritonX and PBS solution at a 1:100 dilution; liver sections were incubated in primary antibodies overnight at 4C. Samples were washed 3x in 0.5% triton X and PBS solution for 30 minutes in between washes before adding secondary antibodies. These antibodies were added to a DAPI with 0.5% tritonX and PBS solution at a 1:250 dilution; liver sections were incubated in secondary antibodies for two hours at room temperature. Samples were washed in 0.5% triton X and PBS solution for 30 minutes before adding RapiClear 1.52 (Sunjin Lab) and incubating the sample overnight. Sections were mounted in Prolong diamond (ThermoFisher) on a superfrost microscope slide and imaged on the Zeiss LM980 Airyscan 2 confocal at 10x-20x magnification. The following primary antibodies were used: anti-mCherry (ab205402), anti-mouse c-KIT (Thermo Fisher 14-1171-82).

### Single-cell RNA sequencing (scRNA-seq)

Mouse or human fetal liver and adult bone marrow cells were prepared as a single cell suspension and prepared for FACS as described above. Single cells were index-sorted into 96-well plates containing lysis buffer as described by Liu et al. Briefly, sorted cells were centrifuged at 4C and snap-frozen on dry ice, and stored at-80C immediately after sorting. Reverse transcription (RT) and cDNA pre-amplification were performed using Smart-seq3 protocol with minor modifications. From the purified cDNA, 1uL was taken for quality control for each well. cDNA concentration and size distribution for each well was determined by Fragment Analyzer (Advanced Analytical). Wells with a concentration less than 1.7ng/uL were excluded his cutoff was determined by measuring the concentration in blank wells with ERCC but no sorted cell. Wells with cDNA concentration above the cutoff value were consolidated and reformatted to a new 384-well plate using the Mosquito X1 liquid handler (SPT Labtech), normalizing each well’s concentration to a range of 1.7–4.0 ng/μL by dilution with UltraPure water. Normalized cDNA was used to prepare Illumina sequencing libraries. Tagmentation was performed by combining 0.4 μL cDNA with 1.2 μL homebrew Tn5 mix (1 ng/μL Tn5 enzyme, 16 mM Tris-HCl pH 7.6, 16 mM MgCl2, and 8% dimethylformamide in UltraPure water). The reaction was stopped by adding 0.4 μL neutralization buffer (0.1% SDS). Indexing PCR reactions were performed by adding 0.4 μL of 5 μM i5 indexing primer, 0.4 μL of 5 μM i7 indexing primer (Integrated DNA Technologies, custom made 7680-plex unique dual index-primer set), and 1.2 μL KAPA HiFi HotStart ReadyMix. PCR amplification was performed using the following program: 72°C for 3 min, 95°C for 30 sec, 98°C for 10 sec, 67°C for 30 sec, 72°C for 60 sec, repeating from step 3 for 10 cycles. For each 384-well plate, 1 μL from each well was pooled, followed by purification using 0.8X volume of AMPure XP beads. The 384-cell library pool from each plate was analyzed for concentration and size distribution using a Fragment Analyzer. Twenty 384-cell library pools were normalized for concentration, pooled to form a 7680-plex library pool, purified, and concentrated using 0.8X AMPure beads. The final 7680-plex library pool was sequenced on a NovaSeq 6000 S4 flow cell (Illumina) to obtain ∼1–2 million 2×150 base-pair paired-end reads per cell.

#### Read mapping

Sequences were demultiplexed using bcl2fastq version 2.19.0.316. 3’ adapter sequences were removed from reads using skewer v0.2.2^39^, and aligned to the hg38 genome (Gencode version GRCm38.p6) for human and GRCm39 for mouse with STAR aligner version 2.6.1d using 2-pass mapping^40^. Briefly, as a first pass, reads for every cell were aligned using STAR genome index generated using the Gencode transcript annotation for the human genome (version 34). Mapped splice junctions for each cell from the first-pass mapping were extracted, aggregated together and a new STAR index was created where any newly discovered splice junctions were included in addition to the existing Gencode annotation during genome index generation. The new STAR index with all known and newly identified splice-junctions was then used for second pass read mapping. Parameters used for STAR mapping were adapted from the ENCODE long-mRNA-pipeline (https://github.com/ENCODE-DCC/long-read-rna-pipeline) recommendations, also detailed in the STAR manual. In addition to the ENCODE recommended options we also used the “--quantMode TranscriptomeSAM” option during second-pass mapping to generate a bam file containing a catalog of all reads mapped to the transcriptome. This bam file was used as input to calculate expression levels of either genes (sum total of expression levels of all known transcript variants) or individual transcripts using RSEM version 1.3.3 with settings “--single-cell-prior”, which accounts for the sparse nature of mRNA detection usually prevalent in scRNA-seq.

#### Data preprocessing

Gene count tables were combined with metadata using the Scanpy python package v.1.8.2. We filtered out genes expressed in fewer than 3 cells, as well as cells with fewer than 500 detected genes or 5000 read counts. The data were normalized using size factor normalization so that every cell has 10,000 read counts, log transformed and scaled to a maximum value of 10. Highly variable genes were computed using default parameters. We then performed principal component analysis, computed the neighborhood graph, and clustered the data using the Leiden method.

### Recombinant VCAM1 binding

Single-cell suspensions were prepared from mouse fetal liver and kept on ice prior to the binding assay. Fetal liver cells (1–2 × 10□cells per condition) were resuspended in ice-cold PBS containing 1% FBS and 1 mM MnCl₂, and incubated with recombinant human VCAM-1–hFc (R&D systems cat. #643-VM) (10 µg/mL) for 45 min on ice. After incubation, cells were washed twice with ice-cold PBS containing 1% FBS and 1 mM MnCl₂, then stained with a fluorophore-conjugated anti-human Fc secondary antibody (AF488 or equivalent) for 20 min on ice in the dark. Cells were washed again and resuspended in PBS with 1% FBS and 1 mM EDTA for flow cytometry analysis. Samples were analyzed by flow cytometry, gating on viable single cells and then on Hoxb5⁺ versus Hoxb5⁻ HSC populations based on prior reporter or phenotypic sorting. VCAM-1 binding was quantified as the percentage of anti-hFc AF488⁺ cells and the mean fluorescence intensity (MFI) within each HSC subset.

### *Ex vivo* VLA-4 blockade and transplantation

E14.5 fetal liver *Hoxb5*-mCherry⁺ HSCs were purified by fluorescence-activated cell sorting (FACS) from Hoxb5-tri-mCherry embryos. Sorted Hoxb5-mCherry⁺ HSCs were incubated for 45 minutes at 4°C in staining buffer containing either anti-VLA-4 antibody (bioxcell clone PS/2 cat. #BE0071) (10 µg/mL) or rat IgG2b (bioxcell clone MPC-11 cat. #BE0086)isotype control antibody (10 µg/mL). Following incubation, cells were washed once with PBS containing 2% FBS and transplanted into lethally irradiated CD45.1 recipient mice together with 2 × 10□ whole bone marrow competitor cells. Peripheral blood and bone marrow donor chimerism were analyzed 4 weeks after transplantation by flow cytometry.

### Quantification and statistical analysis

For all experiments, *n* indicates the number of biologically independent replicates and are indicated in the figure legends. All statistical analysis was performed using GraphPad Prism (GraphPad Software) unless otherwise noted. The statistical tests used (parametric or nonparametric *t*-tests) are noted in the figure legends. Error bars denote mean□±□S.E.M.

## Acknowledgements

We thank the members of the Weissman lab for advice and discussions; Linda Quinn and Teja Naik for laboratory management; Tal Raveh for assistance with general grant administration and funding acquisition. Aaron McCarty and Charlene Wang for mouse colony management; Catherine Carswell-Crumpton, Cheng Pan, and Joe Pasillas for FACS support. This work was supported by the NIA R36 award R36-AG090859 to A.B, the Stanford University Dean’s Fellowship and the Walter V. and Idun Berry Postdoctoral Fellowship to E.K-S, the ETIUDA grant from the Polish National Science Centre no.UMO-2019/32/T/NZ3/00624 to M.Z, the TL1DK139565 to A.T.B, the NIH/NCI Outstanding Investigator Award R35-CA220434 to I.L.W, and the Virginia and D.K. Ludwig Fund for Cancer Research to I.L.W.

## References

1. Spangrude, G. J., Heimfeld, S., & Weissman, I. L. Purification and characterization of mouse hematopoietic stem cells. Science. 241(4861), 58–62 (1988).

2. Weissman, I.L. Translating stem and progenitor cell biology to the clinic: barriers and opportunities. Science. 287(5457), 1442–1446 (2000).

3. Seita, J., & Weissman, I. L. Hematopoietic stem cell: self-renewal versus differentiation. Wiley Interdiscip. Rev. Syst. Biol. Med. 2(6), 640–653 (2010).

4. Weissman, I.L., Papaioannou V, Gardner R. Fetal hematopoietic origins of the adult hematolymphoid system. Differentiation of Normal and Neoplastic Hematopoietic Cells. (Cold Spring Harbor, NY: Cold Spring Harbor Laboratory Press), pp. 33–47 (1978).

5. Müller, A. M., Medvinsky, A., Strouboulis, J., Grosveld, F., & Dzierzak, E. Development of hematopoietic stem cell activity in the mouse embryo. Immunity. 1(4), 291–301 (1994).

6. Medvinsky, A., & Dzierzak, E. Definitive hematopoiesis is autonomously initiated by the AGM region. Cell. 86(6), 897–906 (1996).

7. Galloway, J. L., & Zon, L. I. Ontogeny of hematopoiesis: examining the emergence of hematopoietic cells in the vertebrate embryo. Curr Top Dev Biol. 53, 139–158 (2003).

8. Orkin, S. H., & Zon, L. I. Hematopoiesis: an evolving paradigm for stem cell biology. Cell. 132(4), 631–644 (2008).

9. Morrison, S. J., Hemmati, H. D., Wandycz, A. M., & Weissman, I. L. The purification and characterization of fetal liver hematopoietic stem cells. Proc. Natl. Acad. Sci. U.S.A. 92(22), 10302–10306 (1995).

10. Ema, H., & Nakauchi, H. Expansion of hematopoietic stem cells in the developing liver of a mouse embryo. Blood. 95(7), 2284–2288 (2000).

11. Christensen, J. L., Wright, D. E., Wagers, A. J., & Weissman, I. L. Circulation and chemotaxis of fetal hematopoietic stem cells. PLoS biology. 2(3), E75 (2004).

12. Mikkola, H.K., & Orkin, S.H. The journey of developing hematopoietic stem cells. Development (Cambridge, England), 133(19), 3733–3744 (2006).

13. Kikuchi, K., & Kondo, M. Developmental switch of mouse hematopoietic stem cells from fetal to adult type occurs in bone marrow after birth. Proceedings of the National Academy of Sciences of the United States of America, 103(47), 17852–17857 (2006).

14. Göthert, J. R., Gustin, S. E., Hall, M. A., Green, A. R., Göttgens, B., Izon, D. J., & Begley, C. G. In vivo fate-tracing studies using the Scl stem cell enhancer: embryonic hematopoietic stem cells significantly contribute to adult hematopoiesis. Blood, 105(7), 2724–2732 (2005).

15. Bowie, M. B., Kent, D. G., Dykstra, B., McKnight, K. D., McCaffrey, L., Hoodless, P. A., & Eaves, C. J. Identification of a new intrinsically timed developmental checkpoint that reprograms key hematopoietic stem cell properties. Proceedings of the National Academy of Sciences of the United States of America, 104(14), 5878–5882 (2007).

16. Crisan, M., & Dzierzak, E. The many faces of hematopoietic stem cell heterogeneity. Development (Cambridge, England), 143(24), 4571–4581 (2016).

17. Kim, I., He, S., Yilmaz, O. H., Kiel, M. J., & Morrison, S. J. Enhanced purification of fetal liver hematopoietic stem cells using SLAM family receptors. Blood, 108(2), 737–744 (2006).

18. Kiel MJ, Yilmaz OH, Iwashita T, Yilmaz OH, Terhorst C, Morrison SJ. SLAM family receptors distinguish hematopoietic stem and progenitor cells and reveal endothelial niches for stem cells. Cell, 121(7):1109–1121 (2005).

19. Karimzadeh, A., Varady, E. S., Scarfone, V. M., Chao, C., Grathwohl, K., Nguyen, P. U., Ghorbanian, Y., Weissman, I. L., Serwold, T., & Inlay, M. A. Absence of CD11a Expression Identifies Embryonic Hematopoietic Stem Cell Precursors via Competitive Neonatal Transplantation Assay. Frontiers in cell and developmental biology, 9, 734176 (2021).

20. Cheshier, S. H., Morrison, S. J., Liao, X., & Weissman, I. L. In vivo proliferation and cell cycle kinetics of long-term self-renewing hematopoietic stem cells. Proceedings of the National Academy of Sciences of the United States of America, 96(6), 3120–3125 (1999).

21. Passegué E, Wagers AJ, Giuriato S, Anderson WC, Weissman IL. Global analysis of proliferation and cell cycle gene expression in the regulation of hematopoietic stem and progenitor cell fates. J Exp Med, 202(11):1599–1611 (2005).

22. Pietras, E. M., Warr, M. R., & Passegué, E. Cell cycle regulation in hematopoietic stem cells. The Journal of cell biology, 195(5), 709–720 (2011).

23. Bowie, M. B., McKnight, K. D., Kent, D. G., McCaffrey, L., Hoodless, P. A., & Eaves, C. J. Hematopoietic stem cells proliferate until after birth and show a reversible phase-specific engraftment defect. The Journal of clinical investigation, 116(10), 2808–2816 (2006).

24. Chen, J. Y., Miyanishi, M., Wang, S. K., Yamazaki, S., Sinha, R., Kao, K. S., Seita, J., Sahoo, D., Nakauchi, H., & Weissman, I. L. Hoxb5 marks long-term haematopoietic stem cells and reveals a homogenous perivascular niche. Nature, 530(7589), 223–227 (2016).

25. Potocnik, A. J., Brakebusch, C., & Fässler, R. Fetal and adult hematopoietic stem cells require beta1 integrin function for colonizing fetal liver, spleen, and bone marrow. Immunity, 12(6), 653–663 (2000).

26. Wagers AJ, Allsopp RC, Weissman IL. Changes in integrin expression are associated with altered homing properties of Lin(-/lo)Thy1.1(lo)Sca-1(+)c-kit(+) hematopoietic stem cells following mobilization by cyclophosphamide/granulocyte colony-stimulating factor. Exp Hematol, 30(2):176–185 (2002).

27. Wagers AJ, Weissman IL. Differential expression of alpha2 integrin separates long-term and short-term reconstituting Lin-/loThy1.1(lo)c-kit+ Sca-1+ hematopoietic stem cells. Stem Cells, 24(4):1087–1094 (2006)

28. Qian H, Georges-Labouesse E, Nyström A, et al. Distinct roles of integrins alpha6 and alpha4 in homing of fetal liver hematopoietic stem and progenitor cells. Blood, 110(7):2399–2407 (2007).

29. Frenette PS, Subbarao S, Mazo IB, von Andrian UH, Wagner DD. Endothelial selectins and vascular cell adhesion molecule-1 promote hematopoietic progenitor homing to bone marrow. Proc Natl Acad Sci U S A, 95(24):14423–14428 (1998).

30. Xiang J, Almeida L, Pasillas J, et al. Lineage tracing of both quiescent G0 and active Hoxb5+ LT-HSCs that actively contribute to homeostatic mouse hematopoiesis. Proc Natl Acad Sci U S A, 122(49):e2513724122 (2025).

31. Fleming WH, Alpern EJ, Uchida N, Ikuta K, Spangrude GJ, Weissman IL. Functional heterogeneity is associated with the cell cycle status of murine hematopoietic stem cells. J Cell Biol, 122(4):897–902 (1993).

32. Nygren JM, Bryder D, Jacobsen SE. Prolonged cell cycle transit is a defining and developmentally conserved hemopoietic stem cell property. J Immunol,177(1):201–208 (2006).

33. Wilson A, Laurenti E, Oser G, et al. Hematopoietic stem cells reversibly switch from dormancy to self-renewal during homeostasis and repair. Cell,135(6):1118–1129 (2008).

34. Williams DA, Rios M, Stephens C, Patel VP. Fibronectin and VLA-4 in haematopoietic stem cell-microenvironment interactions. Nature, 352(6334):438-441 (1991).

35. Papayannopoulou T, Craddock C, Nakamoto B, Priestley GV, Wolf NS. The VLA4/VCAM-1 adhesion pathway defines contrasting mechanisms of lodgement of transplanted murine hemopoietic progenitors between bone marrow and spleen. Proc Natl Acad Sci USA, 92(21):9647–9651 (1995).

36. Craddock CF, Nakamoto B, Andrews RG, Priestley GV, Papayannopoulou T. Antibodies to VLA4 integrin mobilize long-term repopulating cells and augment cytokine-induced mobilization in primates and mice. Blood, 90(12):4779–4788 (1997).

37. Arroyo AG, Yang JT, Rayburn H, Hynes RO. Differential requirements for alpha4 integrins during fetal and adult hematopoiesis. Cell, 85(7):997–1008 (1996).

38. Arroyo AG, Yang JT, Rayburn H, Hynes RO. Alpha4 integrins regulate the proliferation/differentiation balance of multilineage hematopoietic progenitors in vivo. Immunity,11(5):555–566 (1999).

39. Priestley GV, Scott LM, Ulyanova T, Papayannopoulou T. Lack of alpha4 integrin expression in stem cells restricts competitive function and self-renewal activity. Blood, 107(7):2959–2967 (2006).

40. Khan JA, Mendelson A, Kunisaki Y, et al. Fetal liver hematopoietic stem cell niches associate with portal vessels. Science, 351(6269):176-180 (2016).

41. Zhang CC, Lodish HF. Insulin-like growth factor 2 expressed in a novel fetal liver cell population is a growth factor for hematopoietic stem cells. Blood. 2004;103(7):2513–2521.

42. Mesquita Piexoto M, Soares-da-Silva F, Bonnet V, et al. Spatiotemporal dynamics of fetal liver hematopoietic niches. J Exp Med. 2025;222(2):e20240592.

43. Lee Y, Leslie J, Yang Y, Ding L. Hepatic stellate and endothelial cells maintain hematopoietic stem cells in the developing liver. J Exp Med. 2021;218(3):e20200882.

44. Iwasaki H, Arai F, Kubota Y, Dahl M, Suda T. Endothelial protein C receptor-expressing hematopoietic stem cells reside in the perisinusoidal niche in fetal liver. Blood. 2010;116(4):544–553.

45. Li D, Xue W, Li M, et al. VCAM-1+ macrophages guide the homing of HSPCs to a vascular niche. Nature, 564(7734):119-124 (2018).

46. Pang WW, Price EA, Sahoo D, et al. Human bone marrow hematopoietic stem cells are increased in frequency and myeloid-biased with age. Proc Natl Acad Sci U S A. 108(50):20012–20017 (2011).

47. Yanai H, Beerman I. Proliferation: Driver of HSC aging phenotypes?. Mech Ageing Dev,191:111331 (2020).

48. Kirschner K, Chandra T, Kiselev V, et al. Proliferation Drives Aging-Related Functional Decline in a Subpopulation of the Hematopoietic Stem Cell Compartment. Cell Rep,19(8):1503–1511 (2017)

49. Flach J, Bakker ST, Mohrin M, et al. Replication stress is a potent driver of functional decline in ageing haematopoietic stem cells. Nature, 512(7513):198-202 (2014).

50. Yoder MC, Hiatt K. Engraftment of embryonic hematopoietic cells in conditioned newborn recipients. Blood, 89(6):2176–2183 (1997).

51. Fowler JL, Zheng SL, Nguyen A, et al. Lineage-tracing hematopoietic stem cell origins in vivo to efficiently make human HLF+ HOXA+ hematopoietic progenitors from pluripotent stem cells. Dev Cell, 59(9):1110–1131.e22 (2024).

52. Ng ES, Sarila G, Li JY, et al. Long-term engrafting multilineage hematopoietic cells differentiated from human induced pluripotent stem cells. Nat Biotechnol, 43(8):1274–1287 (2025).

