## Supplementary figures and images for "Hoxb5+ fetal liver hematopoietic stem cells establish lifelong hematopoiesis and exhibit enhanced ITGA4-dependent engraftment"

### Supplemental

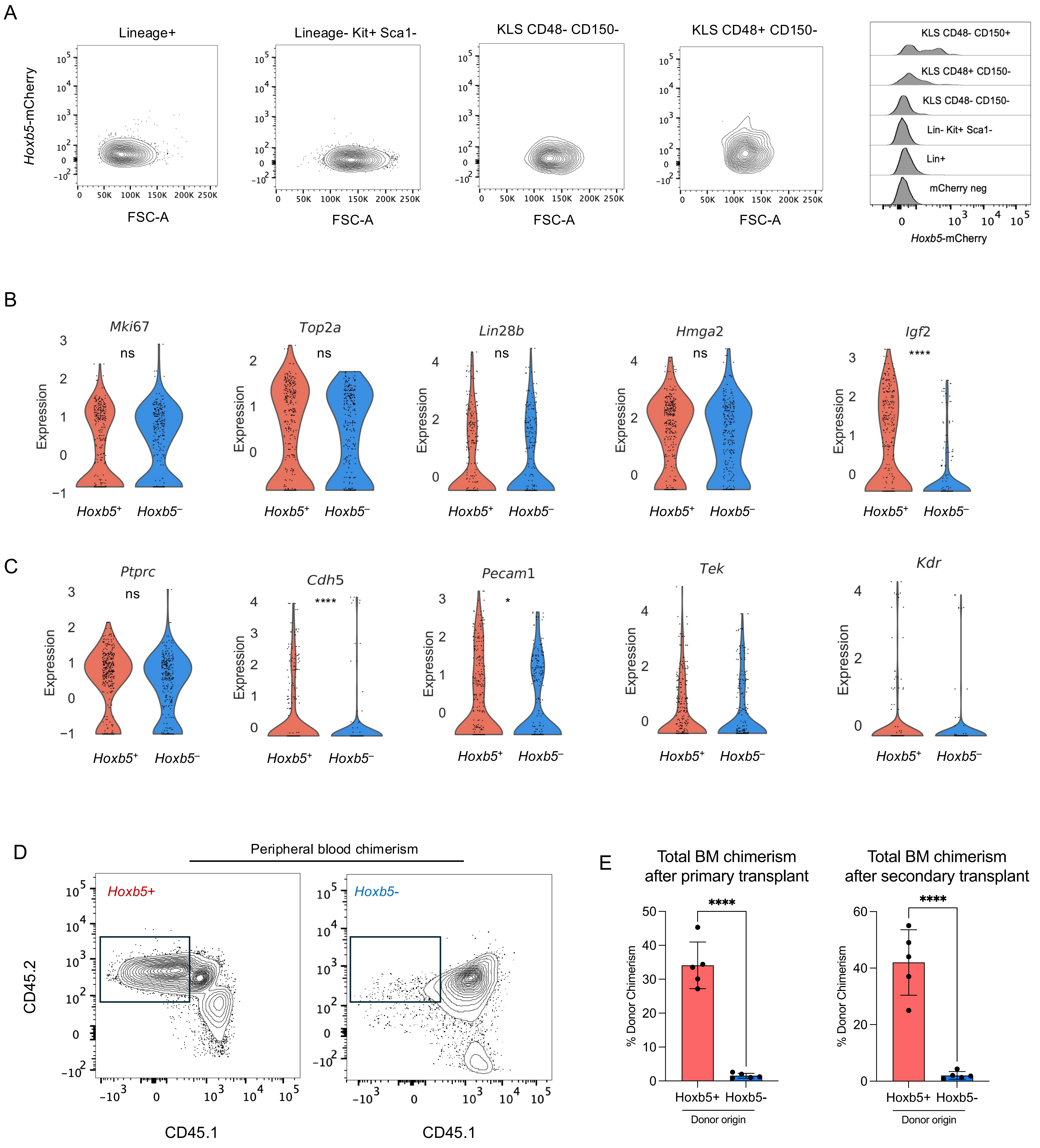

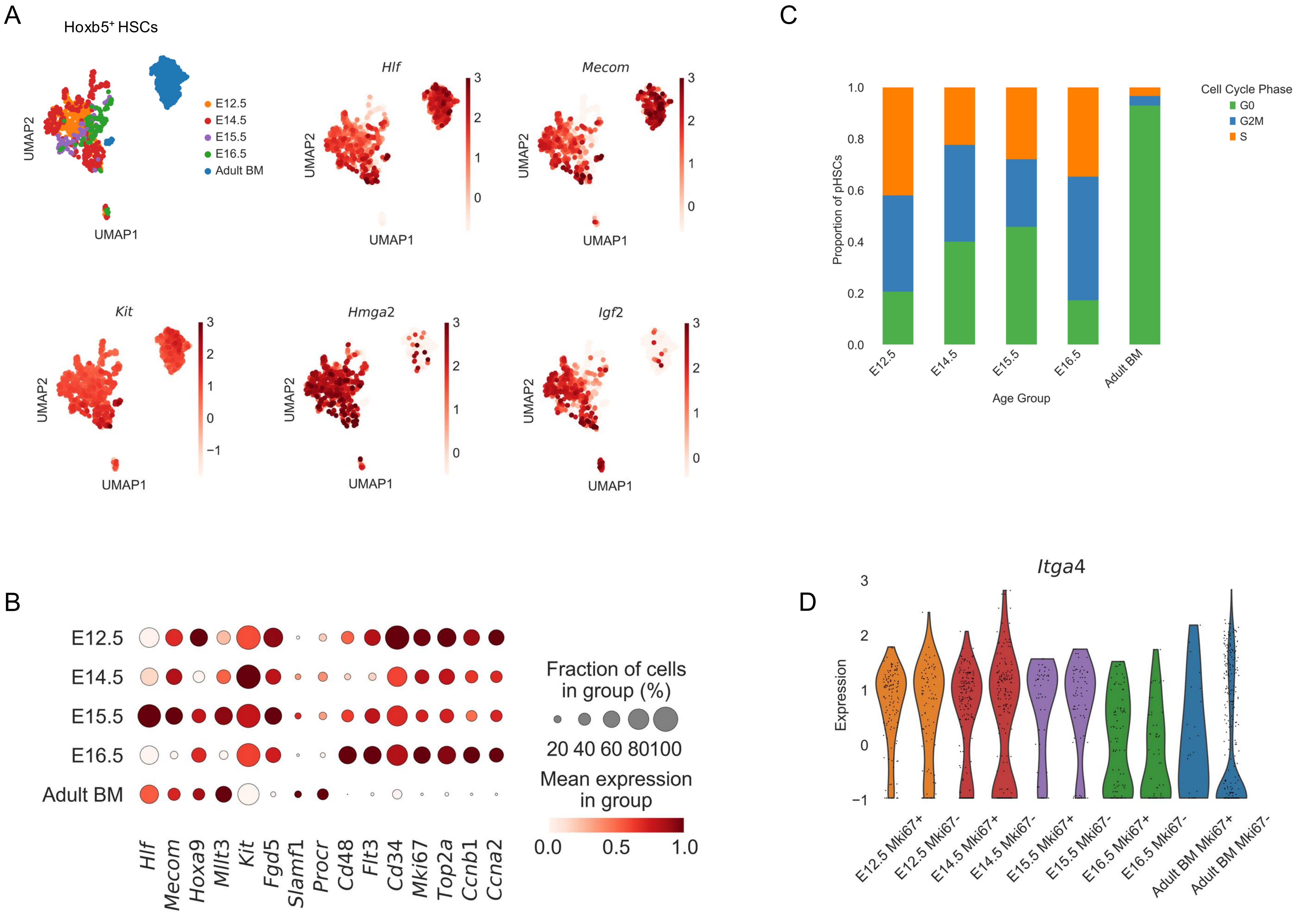

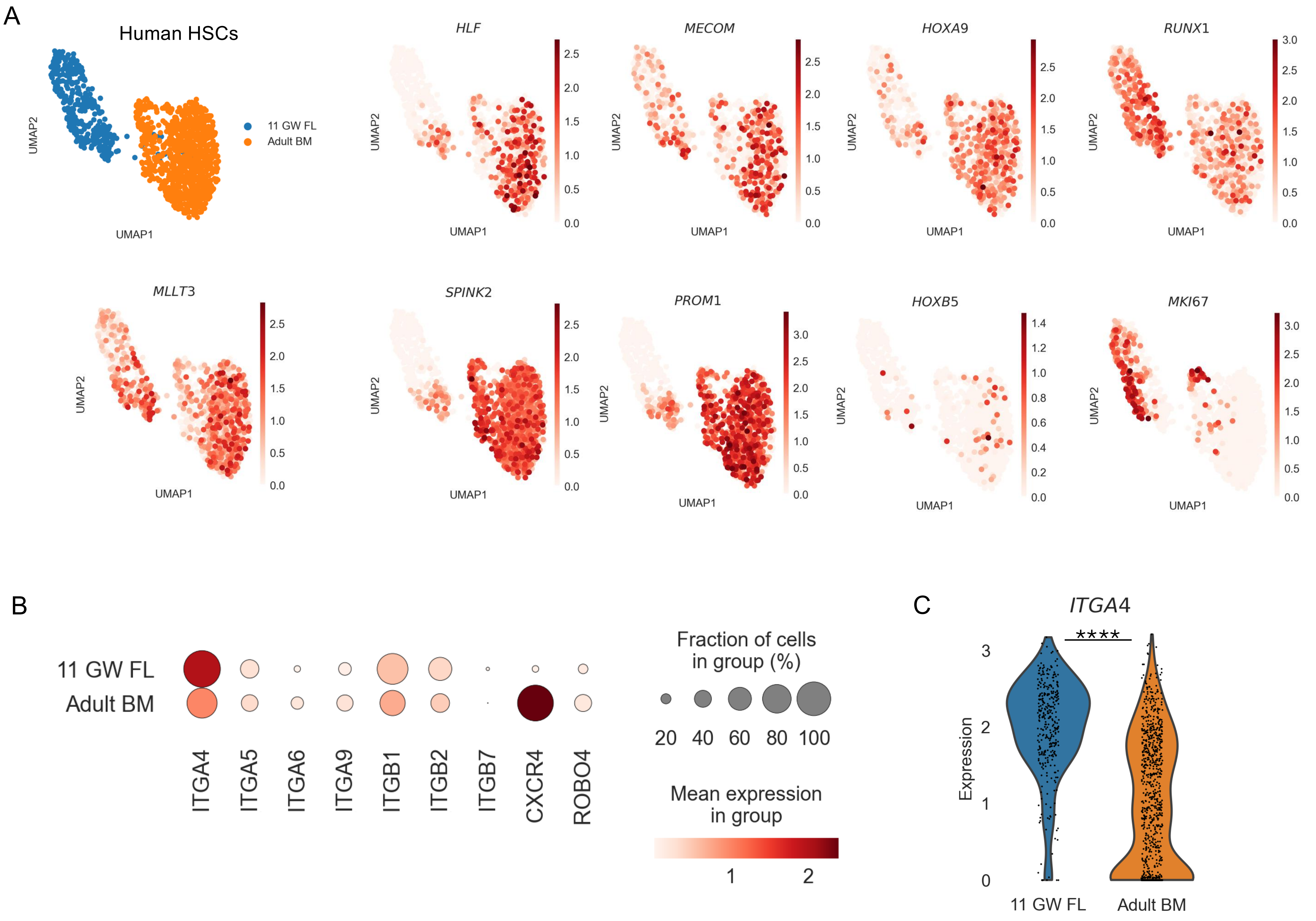

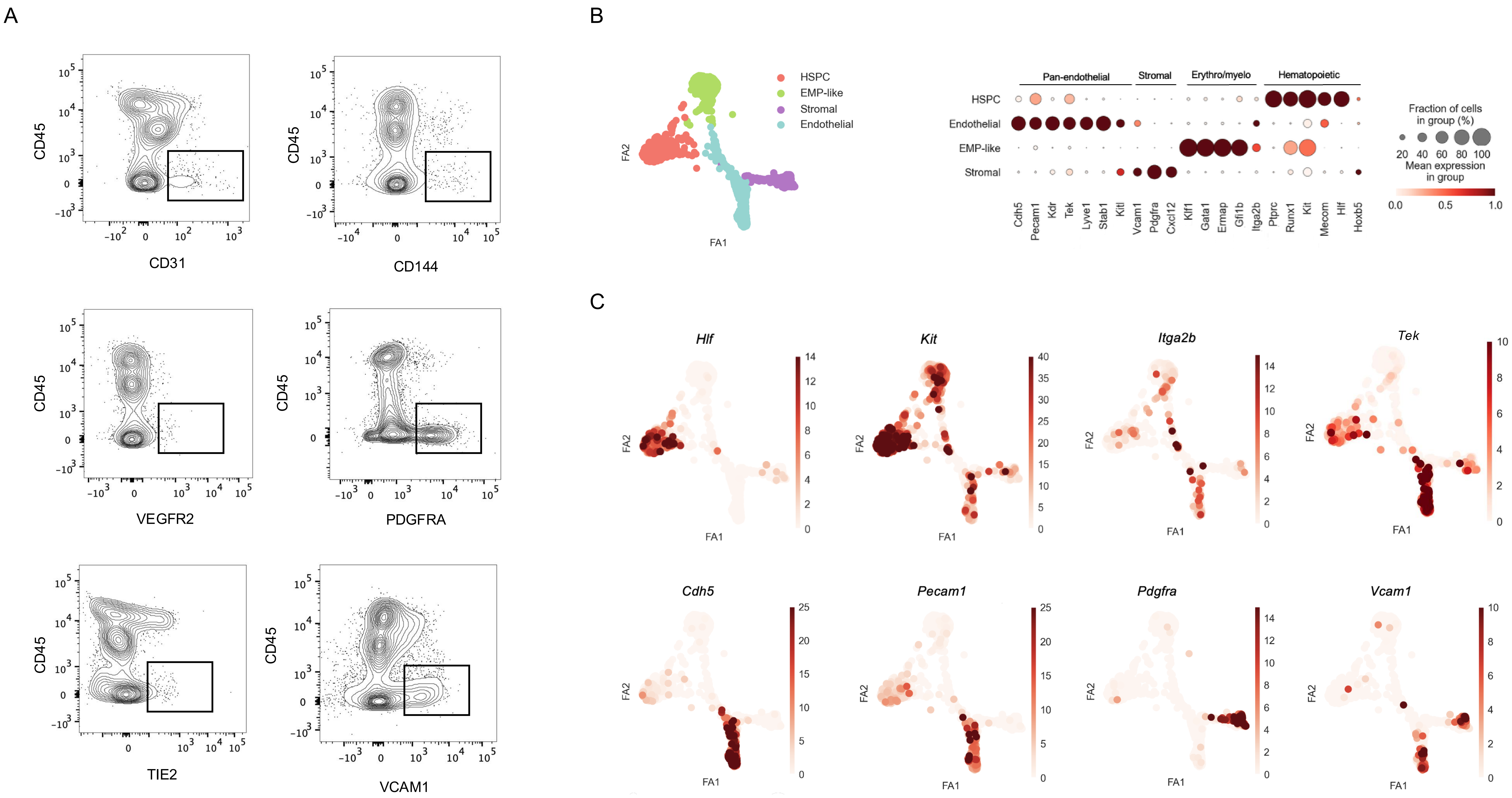
